# Proposed three-phenylalanine motif involved in magnetoreception signaling of an Actinopterygii protein expressed in mammalian cells

**DOI:** 10.1101/2022.12.08.519643

**Authors:** Brianna Ricker, Sunayana Mitra, E. Alejandro Castellanos, Connor J. Grady, Galit Pelled, Assaf A. Gilad

**Affiliations:** Department of Chemical Engineering and Materials Sciences, Michigan State University, East Lansing MI, USA; Department of Biomedical Engineering, Michigan State University, East Lansing MI, USA; Department of Radiology, Michigan State University, East Lansing, MI, USA; Department of Mechanical Engineering, Michigan State University, East Lansing, MI, USA

**Keywords:** glycosylphosphatidylinositol (GPI) anchored protein, magnetoreception, fluorescence microscopy, HaloTag, *Kryptopterus vitreolus*, GCaMP6m, phosphatidylinositol-specific phospholipase C (PI-PLC), site directed mutagenesis

## Abstract

Studies at the cellular and molecular level of magnetoreception – sensing and responding to magnetic fields – is a relatively new research area. As it appears that different mechanisms of magnetoreception in animals evolved from different origins, many questions about the mechanisms remain left open. Here we present new information regarding the Electromagnetic Perceptive Gene (EPG) from *Kryptopterus vitreolus* that may serve as part of the foundation to understanding and applying magnetoreception. Using HaloTag coupled with fluorescent ligands and phosphatidylinositol specific phospholipase C (PI-PLC) we show that EPG is associated to the membrane via glycosylphosphatidylinositol (GPI) anchor. EPG’s function of increasing intracellular calcium was also used to generate an assay using GCaMP6m to observe the function of EPG and to compare its function with homologous proteins. It was also revealed that EPG relies on a motif of three phenylalanine residues in order to function – stably swapping these residues using site directed mutagenesis resulted in a loss of function in EPG. This information not only expands upon our current understanding of magnetoreception but may provide a foundation and template to continue characterizing and discovering more within the field.

**Graphical Abstract:** 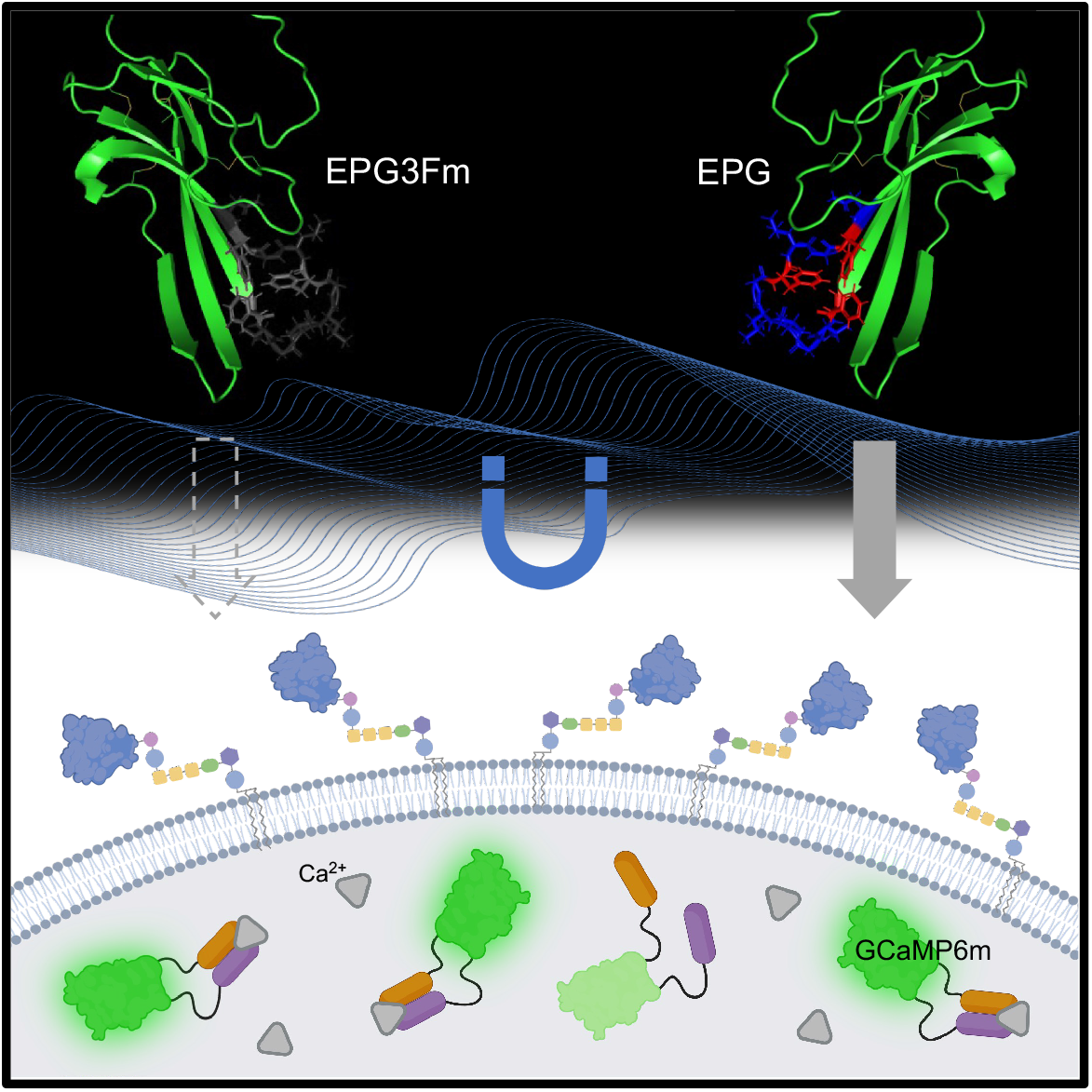

**In Brief:** EPG is a magnetoreceptive GPI anchored protein. Critical to its function is a three-phenylalanine motif which allows EPG to sense and respond to EMF. When expressed in mammalian cell, an increase in intracellular calcium is observed using GCaMP6m. This work represents progress towards understanding magnetoreception for use in future technologies.

**Highlights:**

- EPG is associated to the cell membrane via glycosylphosphatidylinositol anchoring
- In mammalian cells, EPG increases intracellular calcium upon EMF stimulation
- Homologs of EPG from the uPAR/Ly6 family show different responses to EMF
- A three-phenylalanine motif in EPG is critical to its magnetoreceptive ability

## Introduction

In recent years, several organisms from all walks of life have been proposed to have magnetoreceptive properties. Magnetotactic bacteria utilize a membrane-bound crystal containing iron – known as the magnetosome to localize and move in relation to the Earth’s magnetic field^1^. Migratory birds have been proposed to use cryptochromes located in the eye or magnetite-based receptors in the beak to navigate the globe using its inherent magnetic field^2^. Recently, the human brain was demonstrated to be magnetoreceptive^3^. The ability to sense and respond to magnetic fields is well documented in many species and especially in diverse groups of fishes.^4^ Marine animals such as medaka and zebra fish^5^, glass catfish^6^, and eel^7^ represent a large portion of animals that have been proposed to sense magnetic fields and may be used for navigation, predator evasion, or ontogenesis^8^.

Despite the volume of research dedicated to magnetoreception, its exact mechanism remains elusive – especially that which is present in marine life^9^. In an effort to learn more about magnetoreception, the novel Electromagnetic Perceptive Gene (EPG) was cloned from the magnetoreceptive glass catfish *Kryptopterus vitreolus^10^*. EPG has been shown to be functional when expressed in mammalian cells indicated by an increase in intracellular calcium upon stimulation with an electromagnetic field (EMF)^10,11^. This indicated potential for EPG be used to remotely treat nervous system disorders^12^. EPG can potentially be used in synthetic biological technologies as a way of remotely controlling the activation of a system^13^. In order to utilize EPG effectively in these applications, it is essential to characterize its structure and function so it can be optimized for the purpose at hand.

In this project we aimed to elucidate information about the structure, function, localization, and molecular signaling pathway of EPG expressed in mammalian cells. HaloTag proved to be useful throughout the project as a tag that forms strong covalent bonds with various substrates that serve many diverse purposes^14^. HaloTag and its fluorescent ligands were used for imaging purposes that allowed for simple elucidation of EPGs localization, and visualization in cell lysate run on SDS-PAGE. Overall, this study provides important pieces of information about EPG’s structure and function that lead us closer to using magnetoreception as an efficient synthetic tool as well as understanding magnetoreception at the cellular and molecular level.

## Results and Discussion

### EPG is a membrane associated protein with its N terminus located extracellularly

It has been previously reported that EPG was associated to the cell membrane in HEK293 cells.^11^ The next meaningful question to answer is what orientation EPG takes in relation to the membrane. To that purpose, two vectors were created that fuse EPG to HaloTag. Halo-N-EPG consists of HaloTag fused to the N terminus of EPG; preceding HaloTag is EPG’s known signal sequence.^10^ Halo-C-EPG consists of HaloTag fused to the C terminus of EPG. These HaloTag-EPG fusion proteins allow for the use of selectively permeable HaloTag ligands to determine the membranal orientation of EPG. One fluorescent ligand used, Janelia Fluor X 650 (JFX650), is permeable to the cell membrane. The other fluorescent ligand used, Alexa Fluor 488 (AF488), is impermeable to the cell membrane.

HeLa cells expressing Halo-N-EPG were labelled with JFX650 (Fig. 1A) and AF488 (Fig. 1B). Binding of JFX650 indicates adequate expression of Halo-N-EPG by the cells, while binding of AF488 indicates that, minimally, the N terminus of EPG is exposed to the extracellular space. HeLa cells expressing Halo-C-EPG were labelled with JFX650 (Fig. 1C) and AF488 (Fig. 1D). Binding of JFX650 indicates adequate expression of Halo-C-EPG by the cells, and the lack of binding of AF488 indicates, minimally, that HaloTag (and the associated C terminus of EPG) is not exposed to the extracellular space.

**Figure 1.**
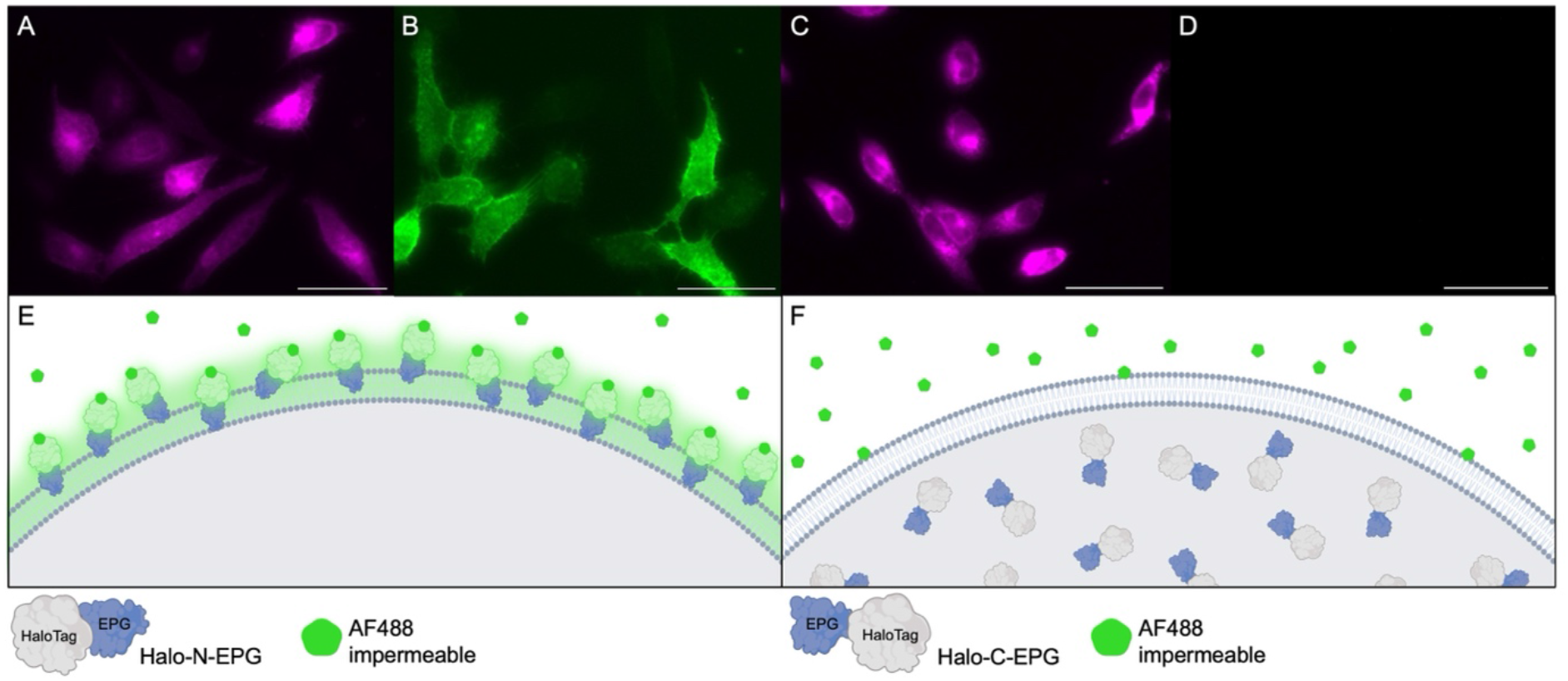
Fluorescently labelled HaloTag indicates EPG’s cellular localization. HeLa cells expressing either Halo-N-EPG (A&B) or Halo-C-EPG (C&D) labelled with fluorescent HaloTag ligands. A&C were labelled with membrane permeable JFX650; indicative of adequate expression of HaloTag-EPG fusion constructs in HeLa cells. B&D were labelled with membrane impermeable AF488. As demonstrated in E, Halo-N-EPG expressing cells were labelled with AF488 indicating its presence as a membrane associated protein with its N terminus exposed to the extracellular space. As demonstrated in F, Halo-C-EPG expressing cells were not labelled with AF488, indicating this construct does not exist at the membrane. Scale bars represent 50μm.

### Evidence that EPG is associated to the membrane via glycosylphosphatidylinositol anchoring

EPG is structurally very similar to members of the Ly6/uPAR family. These proteins are characterized by the presence of a three-finger Ly6/uPAR structural domain formed by disulfide bonds between cysteine residues.^15^ A significant portion of proteins in this family are associated to the membrane via glycosylphosphatidylinositol (GPI) anchor.^15^ Another piece of evidence supporting the idea that EPG is a GPI anchored protein is that Halo-C-EPG does not exist at the membrane as indicated in Fig. 1. One probable conclusion is that HaloTag is blocking a critical signaling domain on the C terminal end of EPG in this construct. This putative signaling domain remains consistent with the way that GPI anchored proteins are formed – translation begins at the N terminus and concludes with the C terminal GPI anchoring sequence being cleaved off and replaced with a GPI anchor in the ER.^16^ Additionally, lysate of HeLa cells expressing Halo-C-EPG and Halo-N-EPG present very differently when run on an SDS-PAGE gel (Fig. 2A). Halo-C-EPG presents exactly as expected – 46 kDa – equivalent to the molecular weight of EPG and HaloTag combined. Halo-N-EPG presents at a higher molecular weight, indicating some sort of posttranslational modification. The increase in molecular weight specifically points to the GPI anchor which has been previously described to bind high quantities of SDS effectively ‘increasing’ the molecular weight of GPI anchored proteins.^17^

**Figure 2.**
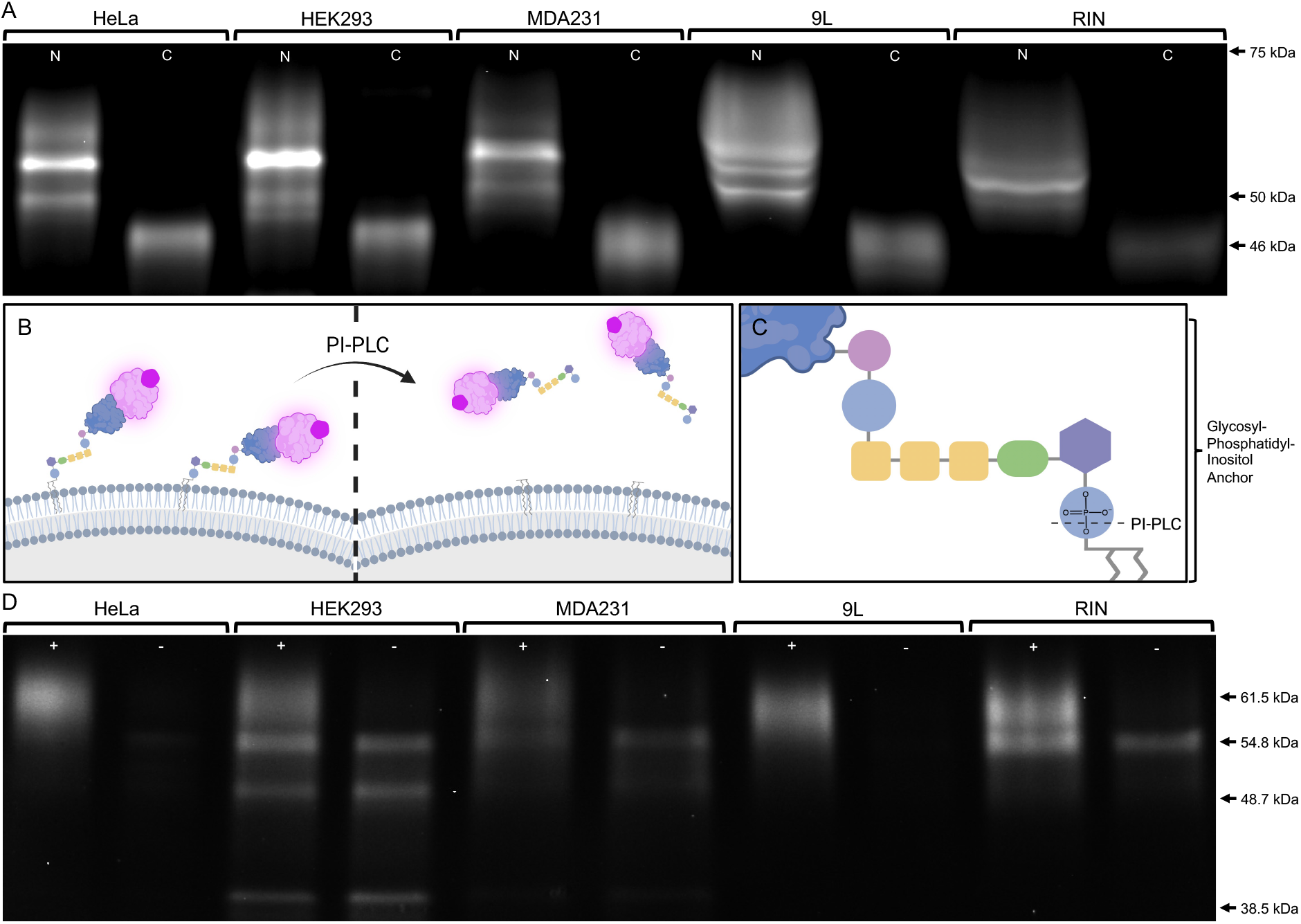
Lysate analysis and PI-PLC digestion signify EPG undergoes posttranslational modification to receive a glycosylphosphatidylinositol anchor. (A) Lysate of various cell lines expressing HaloTag-EPG fusion proteins labelled with JFX650 run on an SDS-PAGE gel visualized with Cy5 exposure. The expected size of HaloTag-EPG fusion proteins is ~46kDa. All cell types expressing Halo-C-EPG (lanes C) exhibited a band at 46kDa. All cell types expressing Halo-N-EPG (lanes N) presented a series of bands between 50-75kDa, indicating potential posttranslational modification allowing Halo-N-EPG to exist at the membrane. (B-D) Various cell lines expressing Halo-N-EPG labelled with JFX650 were treated with PI-PLC which specifically cleaves the phosphatidylinositol of GPI anchors effectively releasing any GPI anchored protein from the membrane as illustrated in B & C. Media from treated (+) and untreated (−) cells visualized by SDS-PAGE with Cy5 exposure shows a band at ~61.5kDa in all treated groups that is not present in the untreated groups—indicative of the presence of a GPI anchor. Different expression levels were observed between the cell lines; lysate and media were diluted to normalize the relative fluorescence of the bands in both A & D.

To test whether EPG is GPI anchored, we utilized the enzyme phosphatidylinositol specific phospholipase c (PI-PLC). As demonstrated in Fig. 2C, this enzyme specifically cleaves the phosphatidylinositol of the GPI anchor. This cleavage effectively releases any GPI anchored protein into the external cellular environment as illustrated in Fig. 2B. Various cell lines expressing Halo-N-EPG were labelled with JFX650 and treated with PI-PLC; control groups were subject to the same treatment without the addition of the PI-PLC. After treatment, media was carefully collected off the top of the cells and run on an SDS-PAGE gel. By using Cy5 exposure, we can specifically visualize HaloTag-EPG fusion constructs within the gel due to the JFX650 ligand. As shown in Fig. 2D, we observe clear bands at 61.5 kDa for the treated groups that are not present in the control groups. Other bands in the gel may be versions of Halo-N-EPG that are misfolded, nonfunctional, or have been excreted for other reasons. As demonstrated in Fig. S1, JFX650 is highly specific to HaloTag and is not likely to bind non-specifically to native proteins in any of the cell lines tested. Fig. S2 shows additional confirmation with the absence of a band in treated cell lysate.

### Analysis of EPG localization and function when HaloTag is fused to each terminus

Previous reports indicate that upon magnetic stimulation, EPG causes an increase in intracellular calcium when expressed in mammalian cells.^10^ We chose to build on this principle, creating a functional assay for EPG. The assay relies on GCaMP6m as a sensor for intracellular calcium levels.^18^ In theory, mammalian cells co-transfected with EPG and GCaMP6m that are magnetically stimulated will experience an influx of cytosolic calcium which will cause GCaMP6m to fluoresce and produce more signal. This idea was confirmed using an EPG-IRES-tdT construct (Fig. 3); HeLa cells were grown in 35 mm tissue culture dishes and stimulated with a custom electromagnetic air-core coil that fits a 35 mm dish in its center.^19^ The coil utilizes double wrapped copper wires that allow for both active and sham stimuli to be produced. The active stimulus is produced when current is run in the same direction in both wires. The sham stimulus is produced when current is run in opposite directions; this anti-parallel configuration cancels the formation of a magnetic field.^19^ The sham is an ideal control in this case because the sample is still subjected to the heat and electricity associated with the magnet, without the magnetic field.^5,20^ Cells were stimulated using a pulse pattern and fluorescence of GCaMP6m was observed for the duration of the experiment using a GFP filter to yield a video such as Video S1. Data for co-transfected cells (determined by overlaying fluorescent images as shown in Figure S3) was gathered by placing regions of interest (ROIs) using the Time Series Analyzer V3 package^21^ for FIJI^22^. The intensity values for each ROI were normalized to the first point in the read – allowing for clearer visual representation of the data. Intensities were then averaged between all ROIs over four experiments to produce the graph shown in Fig. 3G. Cells that received the active stimulus have a higher average intensity than cells that received the sham stimulus and cells that received no stimulus.

**Figure 3.**
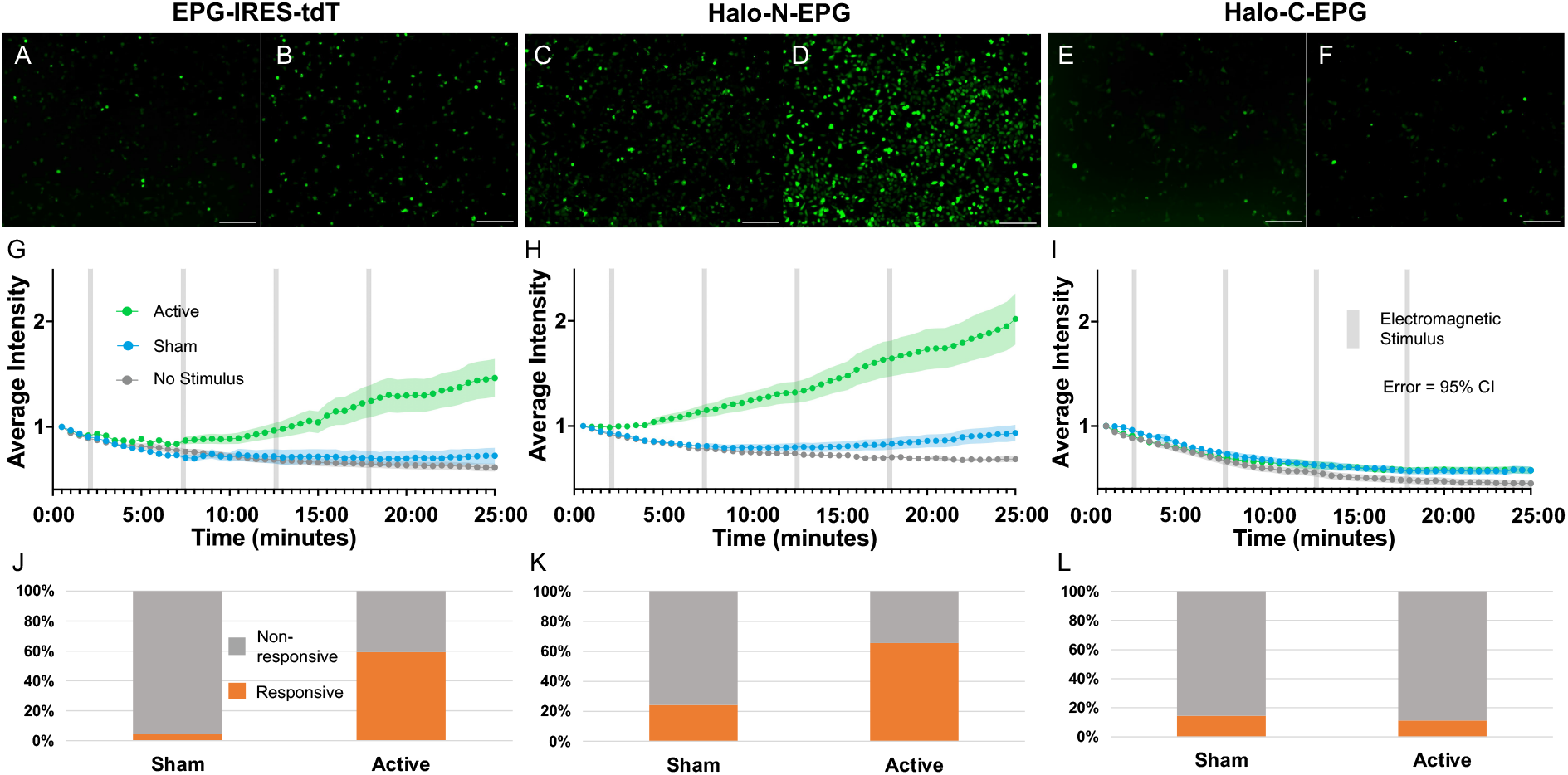
Developing an assay using GCaMP6m to determine if EPG is still functional after the addition of HaloTag. (A-F) HeLa cells expressing GCaMP6m and various EPG-HaloTag fusion constructs visualized before (A, C, E) and after (B, D, F) electromagnetic stimulation. (A-B) Cells expressing EPG-IRES-tdT appear more intense after stimulation. (C-D) Cells expressing Halo-N-EPG appear more intense after stimulation. (E-F) Cells expressing Halo-C-EPG appear relatively unchanged after stimulation. Scale bars represent 200 μm. (G-I) Average intensity of GCaMP6m over time with various electromagnetic stimuli. Active groups received a stimulus of 4.5 A (~14 mT) for 15 s followed by 5 min of rest for 4 pulses (gray bars). Sham groups received a stimulus of 4.5 A (~0.3 mT) with the same pulse pattern as the active group. Control groups were not stimulated. Error bars are representative of 95% CI. Significant increases in intensity were observed between the no stimulus/sham and active groups in both (G) (p<0.0001, Unpaired t test) and (H) (p<0.0001, Unpaired t test). No significant difference was observed between any groups in (I). (J-L) Percentage of individual cells that produced a signal greater than 3 × *SD* + *mean* of the corresponding no stimulus group. Significant differences were observed between sham and active groups in both (J) and (K). No significant difference was observed between sham and active groups in (L). The EPG-IRES-tdT group included n=177, n=134, and n=163 cells over four experiments for no stimulus, sham, and active groups respectively. The Halo-N-EPG group included n=282, n=269, and n=279 cells over four experiments for no stimulus, sham, and active groups respectively. The Halo-C-EPG group included n=160, n=182, and n=189 cells over three experiments for the no stimulus, sham, and active groups respectively.

This functional assay was then applied to the HaloTag-EPG fusion proteins to evaluate if HaloTag altered the function of EPG. Fig. 3H shows that Halo-N-EPG maintained the native function of EPG upon electromagnetic stimulation demonstrated by the increase in average intensity observed in the active group. Fig. 3I shows that Halo-C-EPG does not maintain its function as no increase in intensity was observed. Fig. 3J-K represents data for individual ROIs compared to a threshold of 3 × *SD* + *mean* of their corresponding no stimulus group. ROIs above that threshold were considered ‘responsive’ and cells below the threshold were considered ‘non-responsive’. The EPG-IRES-tdT constructs yielded 59% ‘responsive’ which remains relatively consistent with the Halo-N-EPG groups that yielded 66% ‘responsive’. The Halo-C-EPG groups yielded 11% ‘responsive’ that may be attributed to background cell function.

### Observing how homologs of EPG from different species respond to electromagnetic stimulation

To reinforce the unique function of EPG, we chose to observe how homologs from a range of species would respond to electromagnetic stimulation using the same functional assay. One ideal comparison is a homologous protein from the bluntnose knifefish *Brachyhypopomus gauderio* – a species of electric fish. The protein was identified in the transcriptome of the B.g. fish but remains unnamed and uncharacterized, therefore we will refer to it as ‘B.g.’.^23^ Another ideal homologous control is the protein Bouncer (BNCR) that comes from the zebrafish *Danio rerio* – this species represents a fairly well-studied species of non-electric fish.^24^ Lastly, we chose to compare EPG with a homologous human protein, CD59. This protein represents an ideal control since the experiments are being carried out in human (HeLa) cells. Homology between EPG and the three control proteins is shown in Fig. 4A – conserved amino acids are highlighted in green and similarity of amino acids is indicated by the height and color of the bars on top.

**Figure 4.**
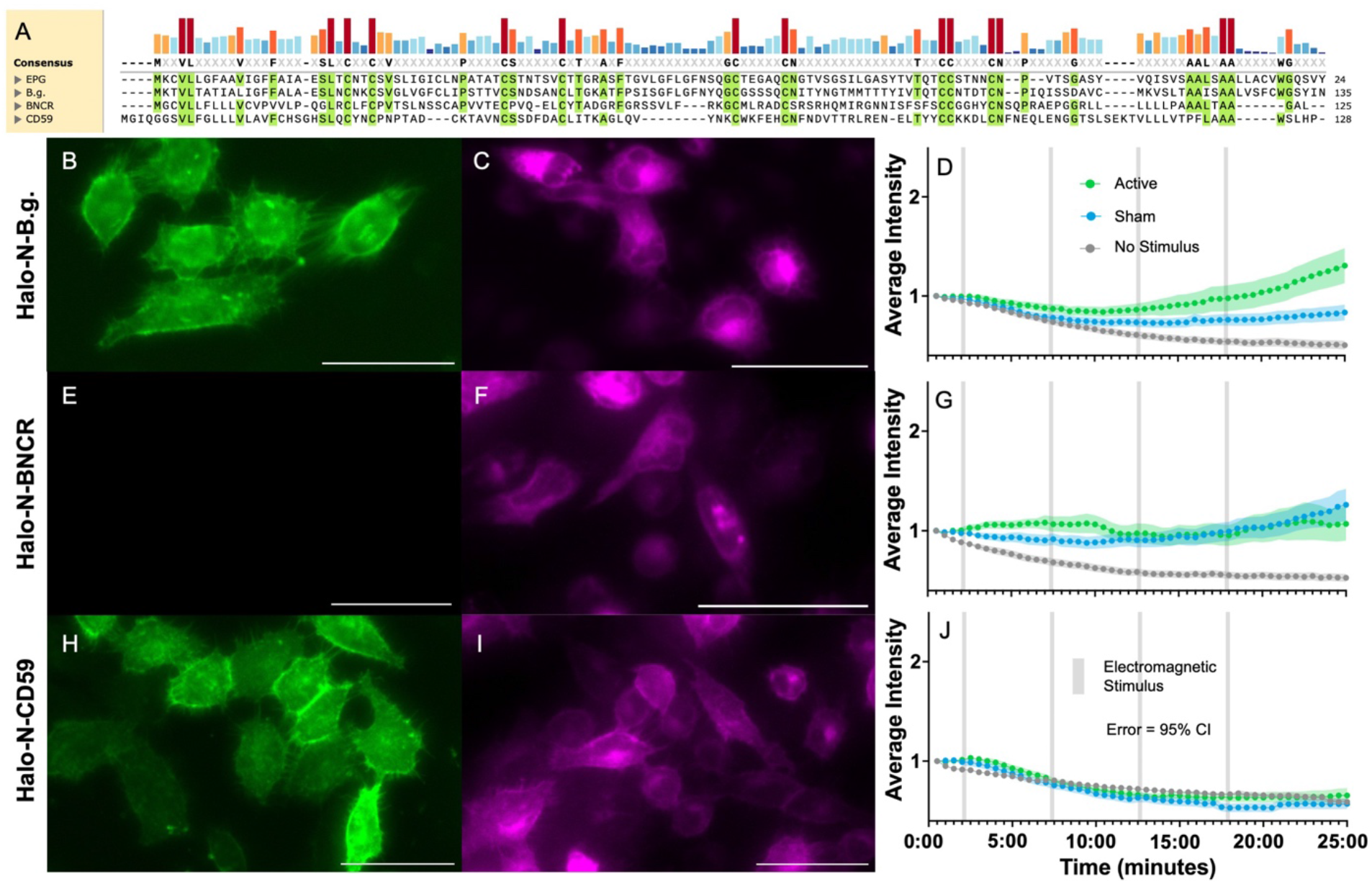
Observing the effect of electromagnetic stimulation on homologs of EPG from different species. (A) Alignment of amino acid sequences for EPG and homologs, B.g., BNCR, and CD59. (B-C) HeLa cells expressing Halo-N-B.g. and labelled with AF488 (B) and JFX650 (C) demonstrating this protein exists at the membrane. (D) Average intensity of GCaMP6m in HeLa cells expressing Halo-N-B.g. with various stimuli. (E-F) HeLa cells expressing Halo-N-BNCR and labelled with AF488 (E) and JFX650 (F) showing the protein, unexpectedly, does not exist at the membrane. (G) Average intensity of GCaMP in HeLa cells also expressing Halo-N-BNCR with various stimuli. (H-I) HeLa cells expressing Halo-N-CD59 labelled with AF488 (H) and JFX650 (I) demonstrating this protein exists at the membrane. (J) Average intensity of GCaMP in HeLa cells also expressing Halo-N-CD59 with various stimuli. (D, G, J) Active groups received a stimulus of 4.5 A which generates a magnetic field of ~14 mT for 15 s followed by 5 min of rest for 4 pulses (gray bars). Sham groups received a stimulus of 4.5 A which generates a magnetic field of ~0.3 mT for 15 s followed by 5 min rest. Control groups were not stimulated. Error bars are representative of 95% CI. All scale bars indicate 50 μm.

B.g. is a similar size to EPG and likely adopts a similar structure to the uPAR/Ly6 family determined by the conserved cysteine residues. Labelling Halo-N-B.g. with impermeable AF488 (Fig. 4B) and permeable JFX650 (Fig. 4C) indicates this protein is adequately expressed in HeLa cells and exists at the membrane. When subject to the functional assay as described above, B.g. seems to have some minimal response to the active and sham stimuli (Fig. 4D). This may be attributed to the protein coming from a species of electric fish; evolutionarily it is possible that the ability to sense electromagnetic fields emerged from the ability to sense electric fields or vice versa.

BNCR is known to adopt the Ly6/uPAR structural domain and is membrane associated – hypothetically via GPI anchor.^24^ Labelling Halo-N-BNCR with impermeable AF488 (Fig. 4E) and permeable JFX650 (Fig. 4F) indicates this particular construct is not associated to the membrane despite being adequately expressed in HeLa cells. In this instance, HaloTag likely interfered with the native structure or function of BNCR. When subject to the functional assay, BNCR may have a minimal response to the sham and active stimuli equally. We hypothesize that because BNCR is involved with egg fertilization^24^, that it may be in some way involved with the hyperpolarization of the egg membrane. Due to the electromagnetic coil design, an electrical field will be present as the voltage coming from the power supply ramps up to and down from the desired voltage. This period lasts about 1 second at the beginning and 1 second at the end of each of the 4 pulses.^19^

CD59 is another member of the Ly6/uPAR family and is ubiquitously expressed in human tissue. It is a GPI anchored protein known to act as an inhibitor of the formation of the membrane attack complex (MAC).^25^ Labelling Halo-N-CD59 with impermeable AF488 (Fig. 4H) and impermeable JFX650 (Fig. 4I) indicate that Halo-N-CD59 does exist at the membrane and is adequately expressed in HeLa cells. When subject to the functional assay, there was no response to any stimuli as shown in Fig. 4J. To make comparison of groups easier, Fig. S4 shows all functional assay graphs side-by-side. Fig. S5 also includes data representing the percentage of individual cells that responded in each group displayed side-by-side for comparison.

### A phenylalanine rich region in EPG is critical for its functionality

To determine how EPG senses and responds to magnetic stimulation, we elected to look closer at its structure in comparison to several homologs from the Ly6/uPAR family. One notable region that stood out is the ‘3F Region’ named for being rich in phenylalanine residues. These residues are relatively conserved between EPG and homologs from species of electric fish, but not homologs from other species. Fig. 5B indicates the position of the three phenylalanine residues in the 3F region in the predicted structure of EPG. The positioning of the aromatic side chains may allow for pi-stacking or facilitate holding a charge. To determine if this region is involved with EPGs function, we knocked out each of the three phenylalanine residues individually and consecutively with the most stabilizing swap determined by ΔΔG calculations which are visually represented in the heatmap shown in Fig. 5A. The first F was swapped for M using site directed mutagenesis and subjected to the functional assay described above. As shown in Fig. 5E the response is relatively consistent with that of native EPG. The second F was swapped in the same manner for W, and its response was also gauged using the functional assay. Fig. 5F demonstrates a loss of function after this mutation. The third F was swapped for W and is demonstrated in Fig. 5G to also cause a loss of function. Finally, FFF was swapped to MWW and tested using the functional assay. We observe again a loss of function.

**Figure 5.**
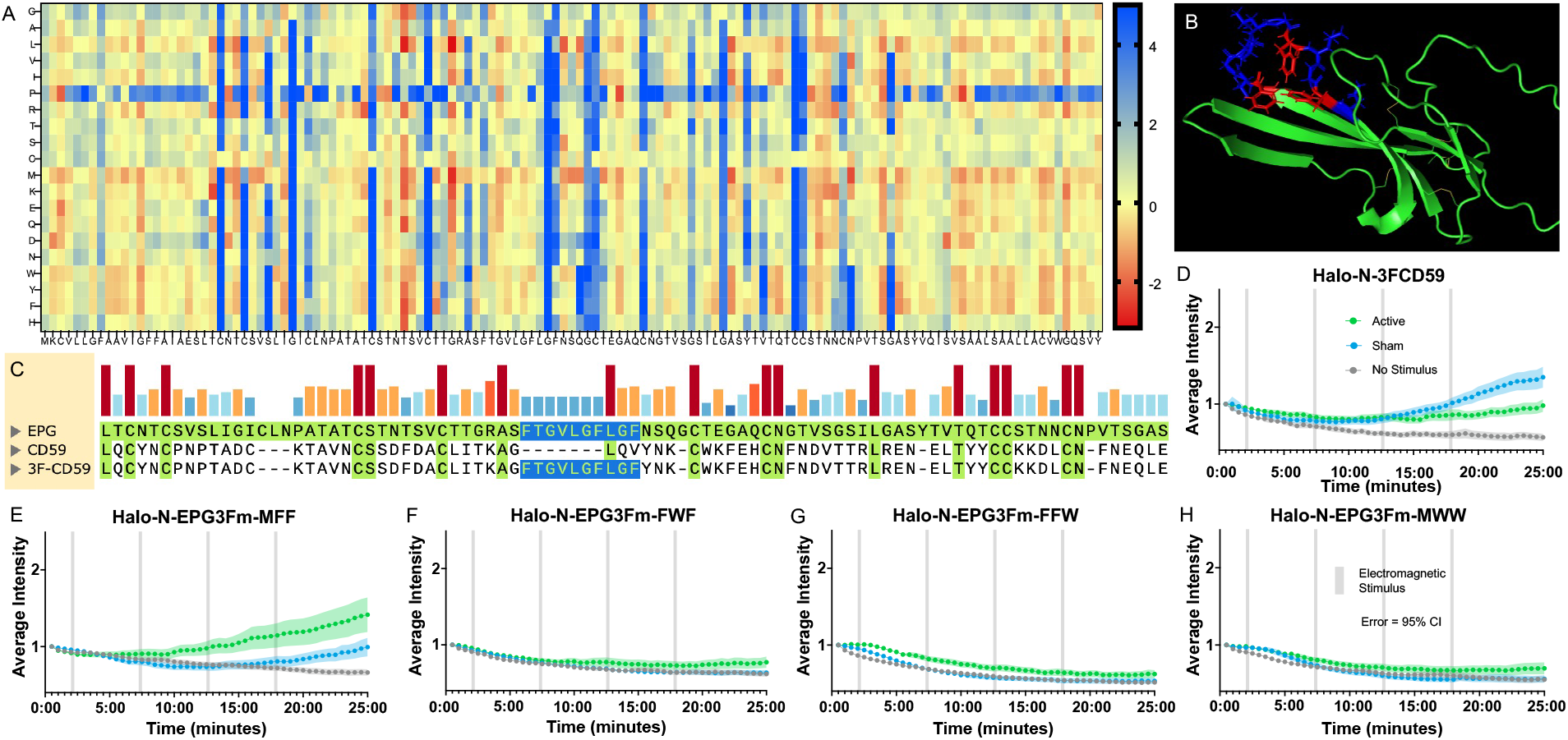
Site directed mutagenesis of the three-phenylalanine region results in change of function. (A) Stability of amino acid swaps for EPG in a heatmap; red is a stabilizing swap, yellow is neutral, and blue is a destabilizing swap. (B) Predicted structure of EPG with the three phenylalanine residues highlighted in red. (C) Alignment of EPG, CD59, and 3FCD59 showing the 3F region (highlighted in blue) from EPG that was inserted into CD59 in a way that conserves the positions of the cysteine residues. (D-H) Average intensity of GCaMP in HeLa cells also expressing various 3F mutants over time with various stimuli. Active groups received a stimulus of 4.5 A (~14 mT) for 15 s followed by 5 min of rest for 4 pulses (gray bars). Sham groups received a stimulus of 4.5 A (~0.3 mT) for 15 s followed by 5 min rest. Control groups were not stimulated. Error bars are representative of 95% CI. (D) Halo-N-3FCD59. (E-H) Versions of EPG with knockouts in the 3F region. Amino acids were swapped with the most stabilizing residue indicated in A. (E) First F in 3F region knocked out. (F) Second F in 3F region knocked out. (G) Third F in 3F region knocked out. (H) All three F residues in 3F region knocked out.

In addition to mutating EPG, we also sought to insert the 3F region into CD59 to determine if we could induce a gain of function in a homolog that previously showed no response. Once again using site directed mutagenesis, the 3F region was inserted into CD59 in a way that conserved the critical positioning of the cysteine residues. Fig. 5C shows the 3F motif highlighted in blue that was taken from EPG and inserted into CD59 to form 3FCD59. When subject to the functional assay (Fig. 5D), 3FCD59 exhibits different functionality to that of CD59 indicating the 3F motif may be essential to the sensing of EMF.

Overall, these findings provide new insight into the structure and function of EPG. This information may serve as a foundation for future work involved with understanding and utilizing the magnetoreceptive abilities of EPG. This work also represents an important step to understanding magnetoreception in biological systems as a whole by providing an example of how a magnetoreceptive protein may function and what its structure may look like. While this study was able to conclude many specifics pertaining to EPG, many questions remain unanswered. Future research may build upon this study by determining the specific pathway EPG takes part in to influence calcium concentrations, or by determining other aspects of EPG’s structure that are critical to its function.

## STAR★Methods

### Resource Availability

#### Lead contact

Further information and requests for resources and reagents should be directed to and will be fulfilled by the lead contact, Dr. Assaf Gilad.

#### Materials availability

Plasmids generated for this study are available upon request.

### Experimental Model and Subject Details

#### Cell Lines

All cell lines were maintained in 25 cm^2^ polystyrene flasks stored in a humidified incubator at 37°C with 5% CO2. HeLa, HEK-293, and MDA-231cells were cultured in DMEM (ThermoFisher) supplemented with 10% FBS (ThermoFisher) and 1% PenStrep (ThermoFisher). 9L/LacZ cells were maintained in DMEM (ThermoFisher) supplemented with 10% FBS (ThermoFisher). RIN-5F cells were cultured in RPMI 1640 medium (ThermoFisher) supplemented with 10% FBS (ThermoFisher) and 1% PenStrep (ThermoFisher).

### Method Details

#### Plasmid Construction and Site Directed Mutagenesis

Primers and g-blocks used to generate constructs for this project were ordered from IDT. The GCaMP6m plasmid was obtained from Addgene.^18^ EPG-IRES-tdT was previously synthesized in the laboratory for use in other projects. Halo-N-EPG was constructed so that the N-terminal signal sequence of EPG preceded the HaloTag sequence followed by the rest of EPG. Halo-C-EPG contains the entirety of EPG followed by the entirety of the HaloTag sequence. All other Halo-N constructs (i.e., BNCR, CD59, B.g.) were cloned by removing the section of EPG following HaloTag and substituting in the gene of interest. EPG’s signal sequence remained preceding HaloTag in all the constructs to increase the likelihood that the proteins would make it to the membrane. All cloning was completed with the NEBuilder HiFi DNA Assembly kit (NEB). To generate the mutant constructs (i.e., Halo-N-3FCD59 and Halo-N-EPG3FK constructs), primers with the desired mutations were used to conduct site directed mutagenesis via PCR.

#### GCaMP6m Functional Assay

HeLa cells were plated in 35mm tissue culture dishes (Falcon) at a density of ~0.1 × 10^6^ cells and allowed to grow for 24 hours. Cells were transfected with both the construct of interest and GCaMP6m using the Lipofectamine 3000 kit (Invitrogen) according to the manufacturer instructions. Cells were labelled with 200 nM JFX650 HaloTag ligand (Janelia Materials) 24 hours post-transfection. The ligand was allowed to bind for 15 minutes, the media was removed followed by two washes with PBS (Corning) to remove excess ligand, and the cells were covered with prewarmed media. Cells were imaged in the BZ-X770 Keyence microscope using a 10X objective while maintained in a Tokai-Hit chamber at 37°C with 5% CO2 and humidity. Cells were visualized with the either the TritC filter to identify tdT or the Cy5 filter to identify cells labelled with JFX650 – both indicative of successful expression of the construct of interest. Cells were visualized with the GFP filter to shown GCaMP6m over 25 minutes. Control groups remained undisturbed for the duration of the experiment. Active and sham groups were stimulated with a custom air-core electromagnetic coil^19^ at 4.5 A (14.5 mT active; 0.3 mT sham) for 15 seconds followed by 5 minutes of rest for 4 pulses for the duration of the experiment.

#### Membrane Localization Imaging

HeLa cells were plated in 96-well glass-bottom plates (Costar) and allowed to grow for 24 hours prior to transfection. Cells were transfected with the construct of interest using the Lipofectamine 3000 kit (ThermoFisher) according to the manufacturer instructions. Half of the wells were labelled with JFX650 (Janelia Materials) and the other half with AF488 (Promega). HaloTag ligands were allowed to incubate with the cells for 15 minutes, excess ligand was then removed by aspirating the media and washing twice with PBS. The cells were covered in prewarmed Fluorobrite DMEM (ThermoFisher) and visualized using the BioTek Cytation 5 Imaging Reader and a 40X objective. AF488 was viewed with the GFP filter and JFX650 was visualized with the Cy5 filter.

#### Lysate Analysis

HeLa, MDA-231, and 9L/lacZ cells were plated in 6-well plates (Corning) at a density of ~0.1 × 10^6^ cells per well, HEK-293 cells were plated in 6-well plates at a density of ~0.3 × 10^6^ cells per well, and RIN-5F cells were plated in 6-well plates at a density of ~0.9 × 10^6^ cells per well and allowed to grow for 24 hours such that all cell lines would be evenly confluent at the time of the experiment. One well of each cell type was transfected with Halo-N-EPG and the other well was transfected with Halo-C-EPG using the Lipofectamine 3000 kit according to the manufacturer instructions. The cells were allowed to grow for 24 hours and were then labelled with 200nM JFX650 (Janelia Materials) and allowed to incubate for 15 minutes. Excess HaloTag ligand was removed by aspirating the media and washing twice with PBS. Each well was lysed in 100 uL of 1X Laemmli sample buffer (Sigma Aldrich) and mechanically separated from the bottom of the plate. The cell lysates were boiled for 5 minutes at 95°C, then 10 uL of each was loaded into a Stain-Free Any kD Mini PROTEAN SDS-PAGE gel (Bio-Rad) and run at 150 V for ~1 hour. The gel was visualized using the Chemi-Doc MP Imaging System (BioRad) with Cy5 exposure to show only JFX650 labelled products. Relative intensity values of bands from this gel were used to ‘normalize’ the amount of sample loaded onto a second gel so that each band would appear with approximately the same intensity. The second gel was run with the varied amount of sample at 100 V for ~1.5 hours and again visualized with Cy5 exposure on the Chemi-Doc.

#### PI-PLC Assay

HeLa, MDA-231, and 9L/lacZ cells were plated in 6-well plates (Costar) at a density of ~0.05 × 10^6^ cells per well, HEK-293 cells were plated in 6-well plates at a density of ~0.15 × 10^6^ cells per well, and RIN-5F cells were plated in 6-well plates at a density of ~0.45 × 10^6^ cells per well and allowed to grow for 48 hours such that all cell lines would be evenly confluent at the time of the experiment. Plates were treated with Poly-D-Lysine (ThermoFisher) prior to plating the cells to ensure cells did not fall off the plate during the experiment. Cells were transfected with Halo-N-EPG using the Lipofectamine 3000 kit (Invitrogen) according to the manufacturer instructions. After 24 hours the cells were labelled with 200 nM JFX650 (Janelia Materials) and allowed to incubate for 15 minutes. Excess HaloTag ligand was removed by aspirating the media and washing twice with PBS. Cells were covered with 250 uL of cold PBS and one well of each cell type was treated with 0.25 units of phosphatidylinositol-specific phospholipase-c (PI-PLC) (ThermoFisher); the other well of each cell type was left untreated. Plates were rocked at 4°C for 20 minutes and the buffer was collected off the top of the cells for analysis. The cell lysates were boiled for 5 minutes at 95°C, then 10uL of each was loaded into a Stain-Free Any kD Mini PROTEAN SDS-PAGE gel (Bio-Rad). 10uL of the collected buffer was mixed with 10 uL of 2X Laemmli sample buffer (Sigma-Aldrich) and boiled for 5 minutes at 95°C, then 10 uL of each was loaded into an SDS-PAGE gel. Both gels were run at 250 V for ~30 minutes. The gels were visualized using the Chemi-Doc MP Imaging System (Bio-Rad) with Cy5 exposure to show only JFX650 labelled products. Relative intensity values of bands from each gel were used to ‘normalize’ the amount of sample loaded onto another gel so that each band would appear with approximately the same intensity. The next set of gels was run with the varied amount of sample at 100 V for ~1.5 hours and again visualized with Cy5 exposure on the Chemi-Doc.

#### Protein Structure Analysis

The structure of EPG was predicted using RoseTTAFold^26^ protein structure prediction software and visualized with the PyMol molecular visualization system. Amino acid alignments of EPG and homologs were generated using SnapGene. ΔΔG calculations were completed with FoldX.

### Quantification and Statistical Analysis

#### GCaMP6m Functional Assay Data Analysis

Twenty-five minutes videos of cells were split into 50 images (one image every 30 seconds) for analysis. The Time Series Analyzer V3^21^ was used in conjunction with FIJI^22^ to place ROIs around viable cells that were confirmed to be co-transfected (i.e., tdT or JFX650 fluorescence and GCaMP6m fluorescence as demonstrated in Figure S3). The ROI was placed so that the cell remained within the borders in all 50 frames with minimal background inclusion. Intensity values for each ROI at each time point were gathered, then normalized to the first point in the read such that each ROI had a starting intensity of one. Intensity values for every ROI over every experiment were averaged and plotted using PRISM to create graphs showing the average intensity over time. The error bars displayed represent a 95% CI. Unpaired t tests were conducted using PRISM to determine significant differences between groups.

#### GCaMP6m Functional Assay Individual Cell Analysis

Intensity over time for each individual cell/ROI was plotted and compared to a threshold of 3 × *SD* + *mean* of the corresponding no stimulus group. Cells were considered ‘responsive’ if they had an intensity greater than the threshold, or ‘non-responsive’ if they had an intensity less than the threshold. Cells were excluded if they oscillated above and below the threshold or exhibited anomalies such as a single spike above the threshold. The total number of ‘responsive’ and ‘non-responsive’ cells was totaled for every experiment to generate percentages shown as bar graphs.

## Author Contributions

Conceptualization of the project was undertaken by B.R., S.M., G.P., and A.A.G. Formal analysis was conducted by B.R. G.P. and A.A.G. were responsible for funding acquisition. B.R., S.M, E.A.C., and C.J.G. developed methodology for the project. A.A.G. was responsible for administration of the project. The manuscript was written by B.R., S.M., and A.A.G.

## Acknowledgments

A.A.G acknowledges financial support from the NIH/NINDS: R01-NS098231; R01-NS104306 NIH/NIBIB: R01-EB031008; R01-EB030565; R01-EB031936; P41-EB024495 and NSF 2027113. The authors thank Dr. Jason Gallant for sharing the B.g. sequence. We also thank Dr. Jens Schmidt, Carlo Barnaba, and David Broadbent for expertise regarding the HaloTag.

