## Supplemental Information for "Proposed three-phenylalanine motif involved in magnetoreception signaling of an Actinopterygii protein expressed in mammalian cells"

### Supporting Information

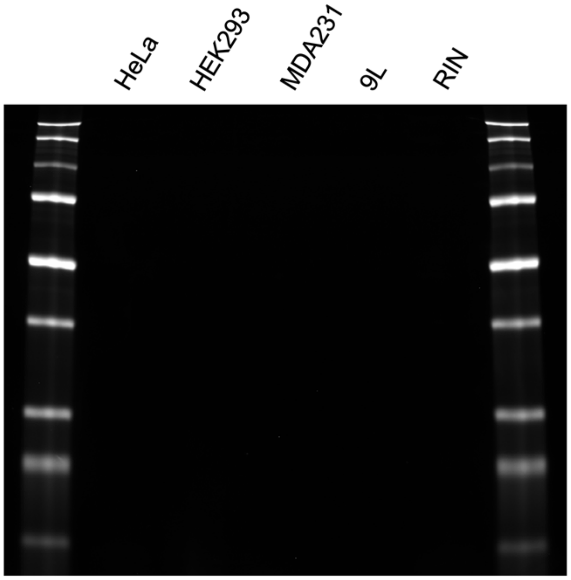


Figure S1. Various non-transfected cell lines labelled with JFX650, lysed and run on an SDS-PAGE gel visualized with Cy5 exposure. The far left and right lanes contain ladders. No bands appear in the lanes containing lysate indicating that the HaloTag ligand JFX650 binds with high specificity to HaloTag and without non-specific binding to proteins native to any of the cell lines tested.

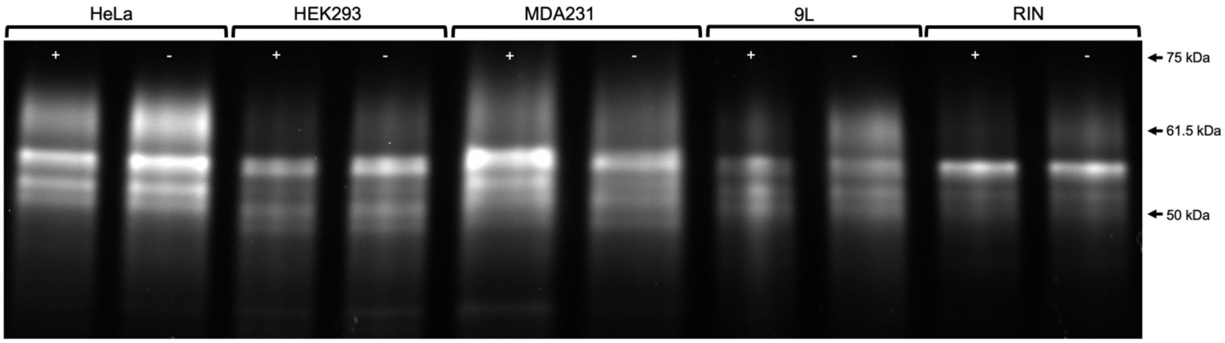


Figure S2. Lysate of cells immediately following PI-PLC digestion. Various cell lines expressing Halo-N-EPG and labelled with JFX650 were subject to treatment with PI-PLC. Media was collected from the top of cells after treatment to collect any GPI anchored proteins that had been cleaved by the PI-PLC. Cell were then immediately washed and lysed. Lysate was run on an SDS-PAGE gel and visualized with Cy5 exposure. We can observe a slightly less prominent band at 61.5 kDa in all cells that were treated (+) versus cells that were not treated (-). This lacking band corresponds to the bands in Figure 2D. Lysate was diluted to normalize the relative fluorescence of the bands.

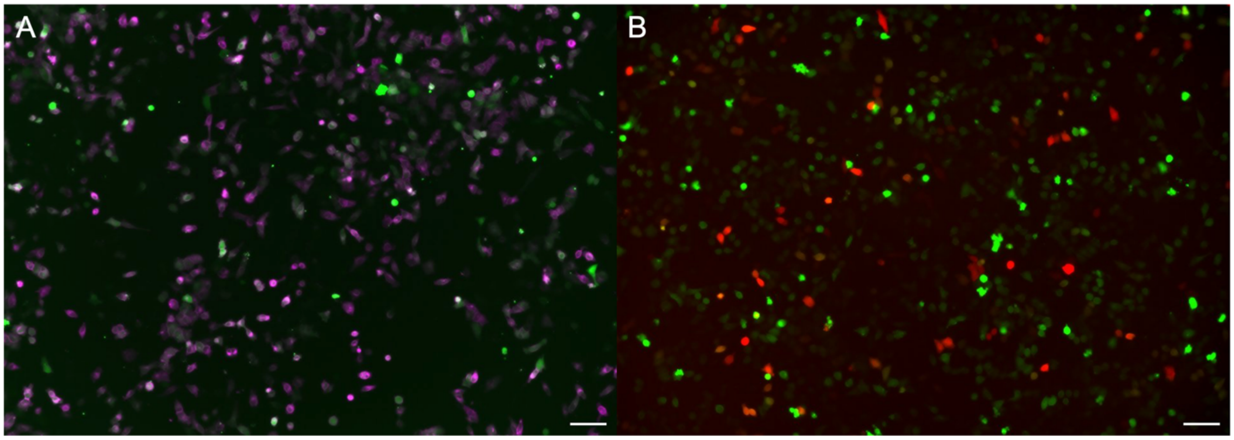


Figure S3. Fluorescent microscopy to identify co-expressing cells. (A) HeLa cells expressing Halo-N-EPG labelled with JFX650 (magenta) and GCaMP6m (green) overlayed to show cells that are co-transfected (white). (B) HeLa cells expressing EPG-IRES-tdT (red) and GCaMP6m (green) overlayed to show cells that are co-transfected (yellow). Scale bar indicates 100 μm.

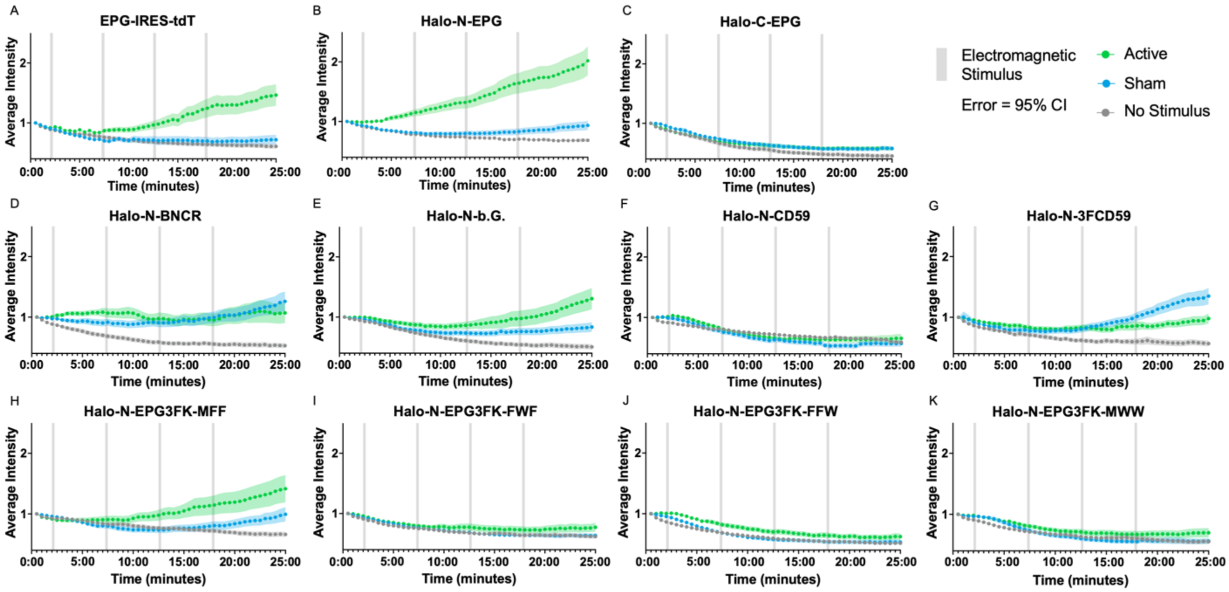


Figure S4. All functional assay graphs displayed next to each other for easier viewing and comparison.

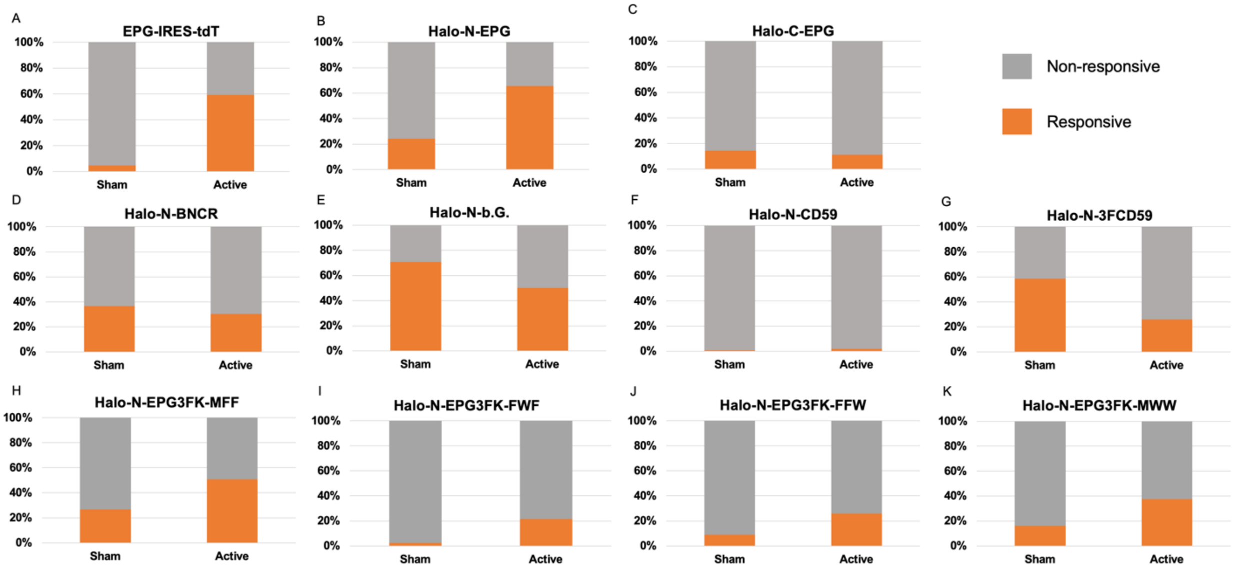


Figure S5. Individual cell analysis from all constructs displayed together for easier viewing and comparison. Percentage of individual cells that produced a signal greater than $3 \times SD + mean$ of the corresponding no stimulus group.
